# MAIT Cells Modulate Innate Immune Cells and Inhibit Colon Cancer Growth

**DOI:** 10.1101/2024.01.16.575894

**Authors:** Olivia J. Cheng, Eric J. Lebish, Owen Jensen, Damian Jacenik, Shubhanshi Trivedi, Jackson Cacioppo, Jeffrey Aubé, Ellen J. Beswick, Daniel T. Leung

## Abstract

Mucosal-associated invariant T (MAIT) cells are innate-like T cells that can be activated by microbial antigens and cytokines and are abundant in mucosal tissues including the colon. MAIT cells have cytotoxic and pro-inflammatory functions and have potentials for use as adoptive cell therapy. However, studies into their anti-cancer activity, including their role in colon cancer, are limited. Using an animal model of colon cancer, we show that peritumoral injection of *in vivo-*expanded MAIT cells into RAG1^-/-^ mice with MC38-derived tumors inhibits tumor growth compared to control. Multiplex cytokine analyses show that tumors from the MAIT cell-treated group have higher expression of markers for eosinophil-activating cytokines, suggesting an association between eosinophil recruitment and tumor inhibition. In a human peripheral leukocyte co-culture model, we show that leukocytes stimulated with MAIT ligand show an increase in eotaxin-1 production and activation of eosinophils, associated with increased cancer cell killing. In conclusion, we show that MAIT cells have a protective role in a murine colon cancer model, associated with modulation of the immune response to cancer, potentially involving eosinophil-associated mechanisms. Our results highlight the potential of MAIT cells for non-donor restricted colon cancer immunotherapy.

**Brief summary:** In models of colon cancer, MAIT cells have anti-tumor activity, associated with increased production of proinflammatory and eosinophil-modulating cytokines.

## Introduction

Mucosal-associated invariant T (MAIT) cells are innate-like T cells that recognize microbial-derived products in a TCR-dependent manner via MR-1, a highly conserved MHC class I-related molecule (1, 2). MAIT cells can be activated with the ligand 5-OP-RU, a microbial riboflavin metabolite, for specific stimulation (3, 4). MAIT cells can also be activated in a TCR-independent manner by cytokines, such as interleukins (IL)-7, IL-12, IL-15, and IL-18 (5–7), broadening the modes of activation and the target cell types. MAIT cells are known for their rapid effector function and cytotoxicity upon activation and have been shown to kill infected target cells (2, 8, 9). Additionally, MAIT cells have the capacity to interact with and modulate other immune cell types, including B cells, neutrophils, and monocytes (10–13). In humans, MAIT cells are abundant in the peripheral blood (14), as well as in mucosal tissues such as the liver (15), lung (16), and gut (17, 18). In mice, MAIT cell frequencies vary by strain and tissue type, but overall have lower frequencies than in humans (19, 20).

Due to the abundance of MAIT cells in various mucosal tissues, interest in MAIT cells has expanded beyond its anti-microbial functions, and recent studies have examined their potential role in mucosal tumors. Reports have suggested both pro- and anti-tumor activity of MAIT cells, with some studies demonstrating a modulating effect on Natural Killer (NK) cell anti-tumor activity (21–23). In addition, due to their lack of alloreactive potential and cytotoxic capacity, MAIT cells have been considered for use in immunotherapy (24–26). Evidently, the role of MAIT cells in cancer is complex and context-dependent, warranting further studies in various types of cancer, particularly their potential to interact with other immune cell types.

As of 2020, human colorectal cancer (CRC) represents about 10% of new cancer cases and death worldwide (27). In addition to surgical resection and adjuvant chemotherapy, the use of immunotherapies is available but limited in efficacy; thus, further understanding of the immune cells involved could help advance therapy (28–30). In human CRC, studies have demonstrated both positive and negative associations between tumor-infiltrating MAIT cells and overall survival (31, 32). MAIT cells accumulate in CRC tumors (17, 31, 33) and tumor-infiltrating MAIT cells display bacterial antigen recognition signatures (34), but the impact of these cells in the tumor microenvironment is not well understood. Tumor-infiltrating MAIT cells retain their cytotoxicity and production of cytokines (17, 35), but have impaired interferon-γ (IFN-γ) production (17, 36) and high levels of exhaustion markers (33, 36). A multi-omics analysis of CRC demonstrated an abundance of intratumor microbes and their immune modulation potential on tumor-infiltrating immune cells, including MAIT cells (32). The prevalence of intratumoral bacteria in CRC (32, 34) and the specificity of MAIT cells for microbial antigens warrant further investigation into the role of MAIT cells in CRC.

Here, we show that peritumoral injection of purified MAIT cells inhibits MC38 cells-derived tumor growth in a mouse model of colon cancer. We show that this tumor inhibition is associated with changes in the immune cell profile and functional changes, including higher levels of myeloid cell-modulating cytokines and chemokines. Culture of human peripheral leukocytes with colon cancer cell line resulted in the activation of eosinophils upon MAIT cell stimulation. Together, our data suggest a novel mechanism by which MAIT cells can modulate innate immunity by affecting eosinophils in a mixed immune cell environment and provide protection in a murine model of colon cancer, highlighting the potential of MAIT cells to be a candidate for CRC immunotherapy.

## Results

### Peritumoral injection of MAIT cells leads to inhibition of tumor growth in the MC38 model of CRC

To test the effect of MAIT cells on colon cancer growth, MAIT cells were expanded in C57BL/6J wildtype (WT) mice for 7 days, isolated (Supplemental Fig. 1), and were peritumorally injected into RAG1^-/-^ mice with MC38 cells-derived tumor 1 day after cancer cell injection (Fig. 1A). We observed that the growth of MC38 cells-derived tumors was significantly lower (p<0.05) when mice were treated with MAIT cells, as shown in the final tumor volume and weight at harvest (Fig. 1B-E). The same effect was also observed when MAIT cells were injected with ligand 5-A-RU and methylglyoxal (MGO) into both MC38-bearing RAG1^-/-^ and C57BL/6J WT mice (Supplemental Fig. 2). These data suggest that MAIT cells provide a protective role in this murine subcutaneous model of colon cancer.

**Figure 1.**
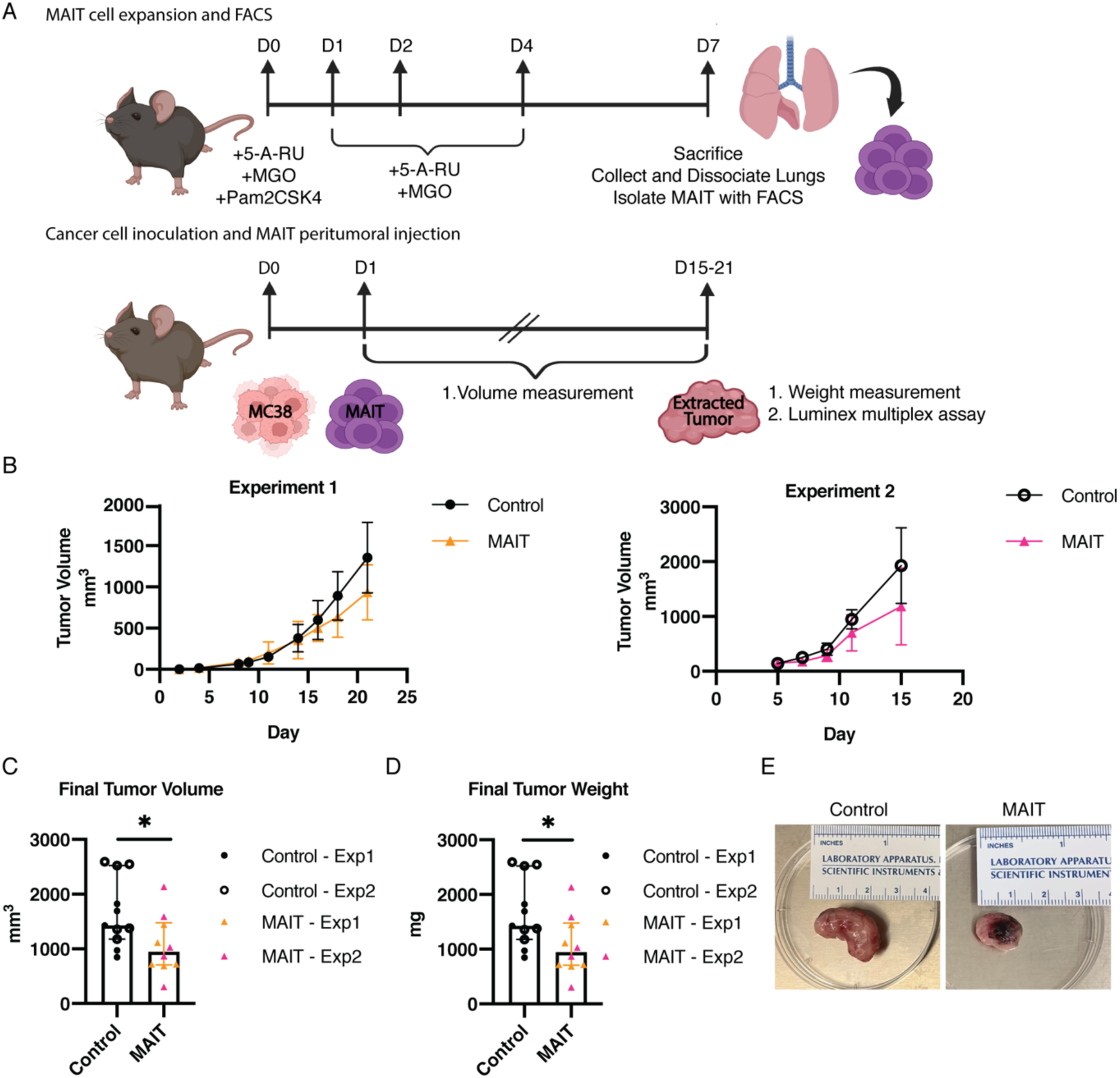
MAIT cell peritumoral injection leads to inhibition of MC38 cells-derived tumor growth. A) Experimental setup of MAIT cell expansion in C57BL/6J mice and MAIT cell peritumoral injection in MC38-bearing RAG1^-/-^ mice. MAIT cells were expanded in C57BL/6J mice for 7 days and sorted MAIT cells were injected peritumorally in RAG1^-/-^ mice 1 day after MC38 cell injection. Control: n = 11 males. MAIT: n = 10 (9 males and 1 female). Tumor volume was monitored during the course of the experiment. Tumors were extracted at the end of the experiment for weight measurement and Luminex multiplex analysis. B) MC38 cells-derived tumor volume in mm^3^ over time from 2 independent experiments. C-D) Tumor measurements. Each dot represents a tumor sample and data is shown as median with interquartile range. C) Volume of extracted tumors in mm^3^ and D) Weight of extracted tumors in mg. *p<0.05 by two-tailed Mann-Whitney U test. E) Representative photos of extracted tumors from each experimental group. Figure created using BioRender.com

### MAIT cell peritumoral injection induces cell death and cytokine production in the tumor microenvironment

Consistent with the observed inhibition of tumor growth (Fig. 1), caspase 3/7 protein activity was detected to be significantly higher in tumors in the MAIT cells-treated group compared to the control group (p<0.01; Fig. 2A), suggesting that the treatment of MC38-bearing mice with MAIT cells leads to increased cell death.

**Figure 2.**
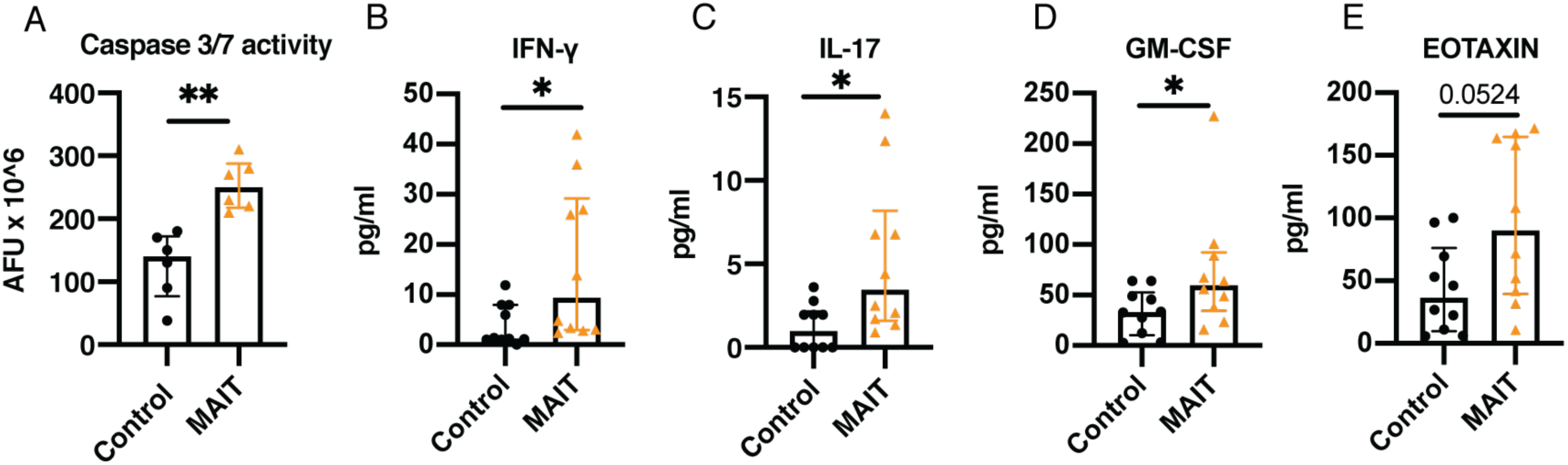
MAIT cell peritumoral injection induces changes in cytokines produced within tumors. Comparison of (A) caspase activity and (B – E) concentration of various cytokines and chemokines released from extracted tumors from Figure 1 between control group and MAIT group. A) Caspase activity is measured in arbitrary fluorescent unit (AFU). B) GM-CSF, C) eotaxin, D) IFN-γ, and E) IL-17. 25 µL of supernatants from 18-hour culture of 8 mg tumor pieces were analyzed using Luminex multiplex assay. Each dot represents an individual sample of supernatant. Data was collected from 2 independent experiments and is shown as median with interquartile range. *p<0.05 **p<0.01 by two-tailed Mann-Whitney U test.

To assess any functional changes in the tumor microenvironment, we analyzed released cytokines and chemokines from the extracted tumor. Tumor pieces (8 mg) were incubated in complete media for 18 hours and the resulting supernatants were analyzed using Luminex multiplex cytokine and chemokine analysis. Out of the 31 cytokines and chemokines analyzed (Supplemental Table 1), we observed a significant increase in the production of various cytokines, including the proinflammatory cytokines IFN-γ, and IL-17, and GM-CSF (p<0.05; Fig. 2B, C, and D), which provides further evidence that the introduction of MAIT cells into MC38 cells-derived tumors leads to the reprogramming of the tumor microenvironment. Additionally, a higher level of eotaxin-1 (p = 0.0524) was detected in the tumors extracted from mice in the MAIT treatment group (Fig. 2E). To determine the plausibility that these cytokines could be produced by activated MAIT cells, we cultured sorted mouse MAIT cells alone *in vitro* with a T cell activation cocktail. We confirmed significantly higher levels of each of the above cytokines compared to unstimulated conditions (Supplemental Fig. 3). Together, we observed functional changes of immune cells and reprogramming of the tumor microenvironment associated with *in vivo* tumor inhibition.

### Activation of human MAIT cells with a MAIT ligand enhances the killing of colon cancer cells

To model the effect of human MAIT cells on colon cancer cells in the presence of other immune cells, we utilized human whole leukocytes co-cultured with the human colon cancer cell line COLO 205 (Fig. 3A). Calcein release assay was used to examine cytotoxicity and culturing with human whole leukocytes for 4 hours leads to specific lysis of COLO 205 cells (Fig. 3B). Activating MAIT cells with the MAIT ligand, 5-A-RU and MGO, significantly increased the specific lysis capacity of the leukocytes (p<0.05; Fig. 3B). Flow cytometry analysis also demonstrates enhanced killing of COLO 205 by whole leukocytes with the addition of the MAIT ligand in a 16-hour co-culture, with a significant reduction in the frequency of CFSE-labeled cells (p<0.05; Fig. 3C). Together, these data show that MAIT cell-specific activation with 5-A-RU enhances the killing of colon cancer cells.

**Figure 3.**
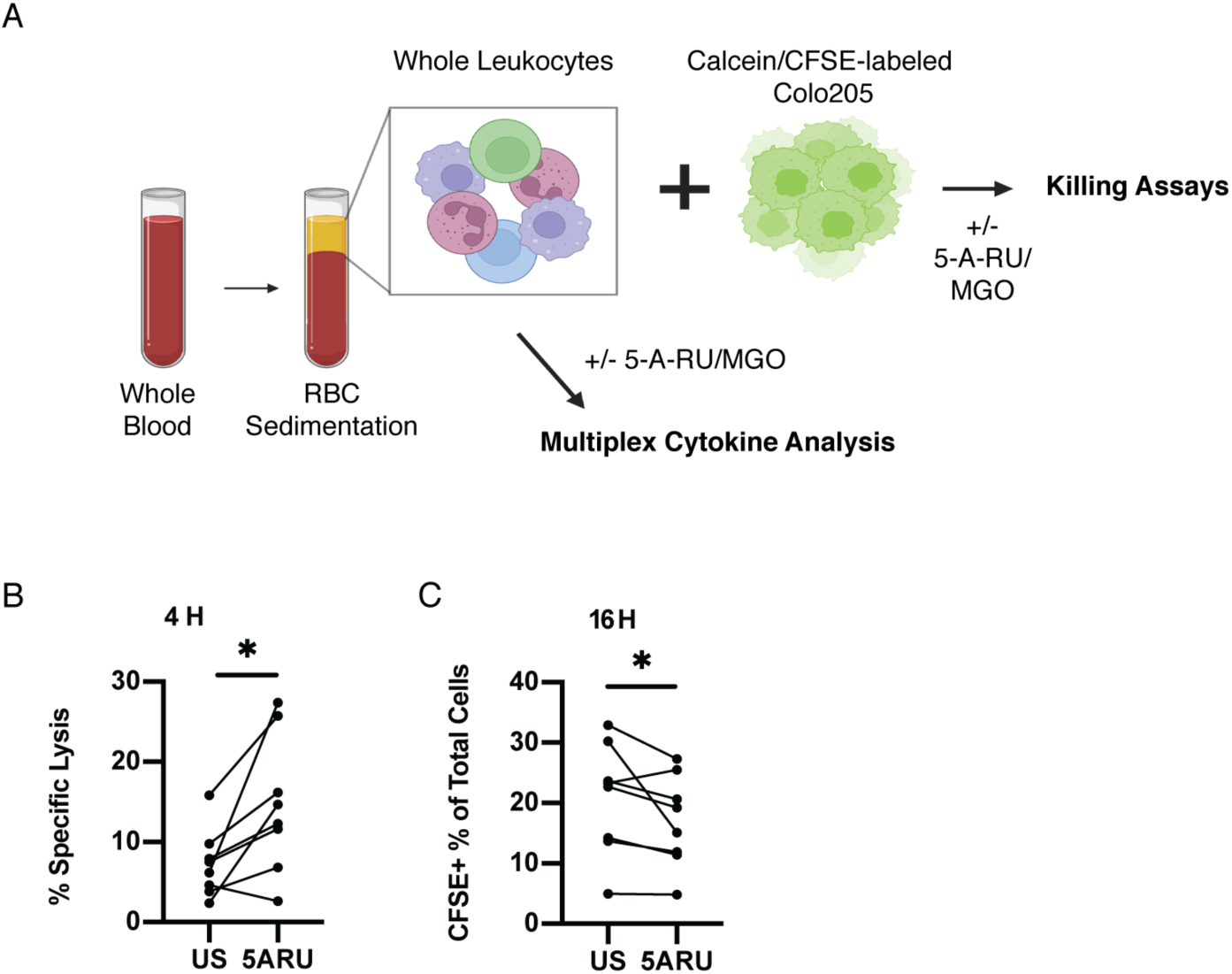
Activation of MAIT cells enhance killing against colon cancer cells. A) Experimental setup of human whole leukocyte isolation and killing assay co-culture with COLO 205. B) Percent specific lysis of COLO 205 co-cultured with human whole leukocytes with or without 5-A-RU/MGO stimulation using a 4-hour Calcein Release Assay. C) Frequency of CFSE+ cells after overnight co-culture of human whole leukocytes with COLO 205 with or without 5-A-RU/MGO stimulation for 16 hours by flow cytometry. Each dot represents a sample from an individual healthy donor and data was collected from 2 independent experiments. Figure created using BioRender.com. *p<0.05 by Wilcoxon ranked test.

### Stimulation of MAIT cells in human peripheral leukocytes induces eotaxin-1 production in MAIT cells and activation of eosinophils in an MR-1 dependent manner

To determine the immune modulating potential of human MAIT cell in a mixed cell population, we used human whole leukocyte cultures to examine the effect of MAIT cell activation in a cell culture model of CRC. MAIT cells were selectively stimulated with 5-A-RU overnight and culture supernatant was analyzed using multiplex cytokine and chemokine assay. In 5-A-RU stimulated cultures, we found a higher level of Macrophage inflammatory protein (MIP)-1α, MIP-1β, and Monocyte chemoattractant protein (MCP)-1 (p< 0.05 and p<0.01; Fig. 4A), suggesting a capacity of MAIT cells to modulate macrophages and monocytes and granulocytes (37–40). Consistent with increase in eotaxin-1 with MAIT treatment in our murine model (Fig 2A), we found that concentrations of IL-5, GM-CSF, and eotaxin-1 levels are significantly higher in cultures stimulated with the MAIT ligand 5-A-RU/MGO compared to unstimulated leukocytes (p<0.05 and p<0.01; Fig. 4A), highlighting the capacity of MAIT cell activation on modulating eosinophils. In addition, we observed that other cytokines known to activate and attract eosinophils, including IL-4 (p<0.01) and IL-8 (p=0.0547) (41, 42), are also significantly higher in 5-A-RU/MGO stimulated cultures compared to control (Fig. 4A). Together, these data demonstrate the immune modulating capacity of human MAIT cell activation in a mixed cell population, specifically the potential to modulate eosinophils.

**Figure 4.**
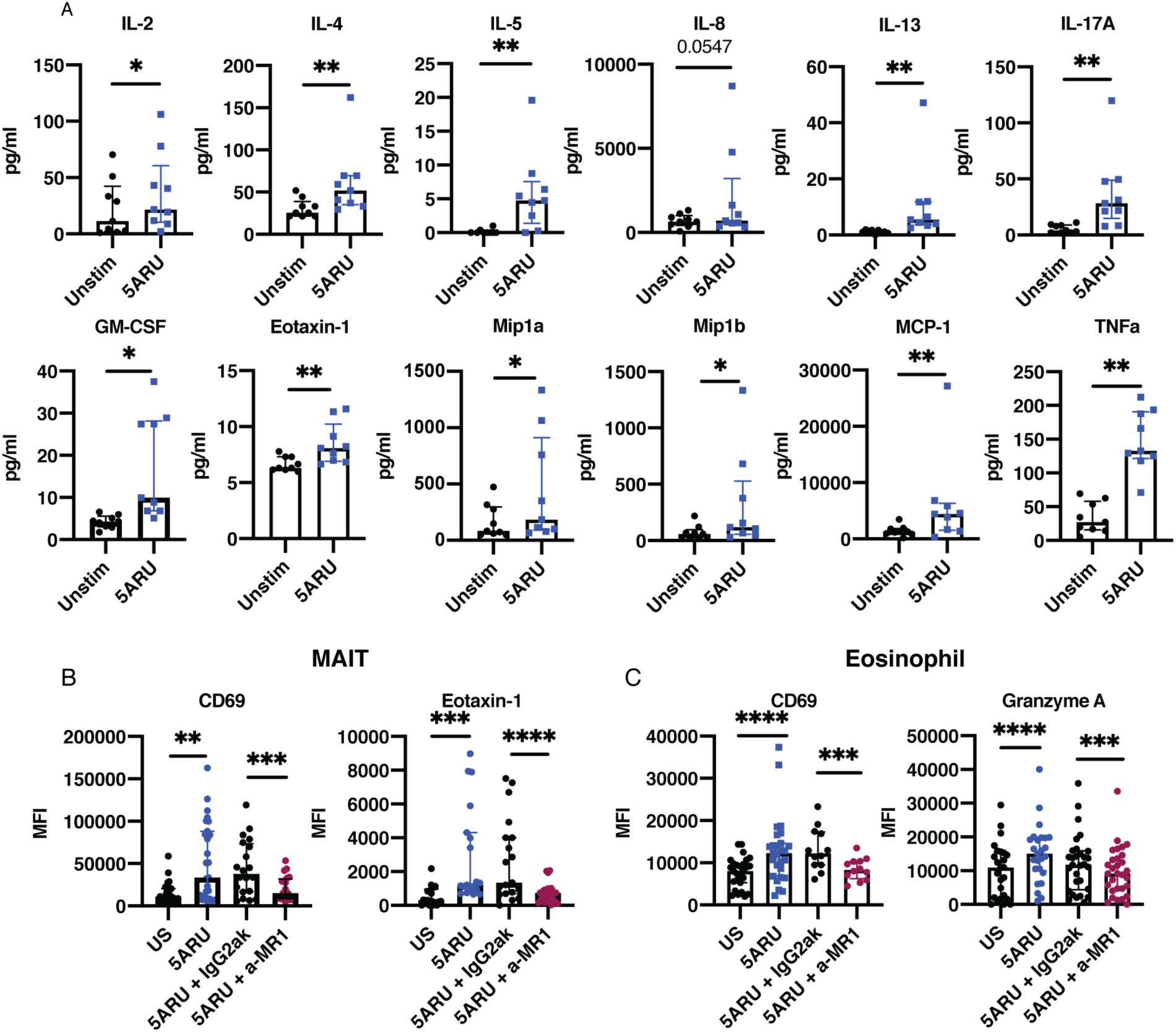
Stimulation of human peripheral leukocytes with MAIT ligand induces eotaxin-1 production in MAIT cells and activation of eosinophils in an MR-1 dependent manner. A) Supernatant concentration of various cytokines and chemokines detected in overnight cultures of human whole leukocytes with or without 5-A-RU/MGO stimulation analyzed by Luminex multiplex assay. Each dot represents an individual sample of supernatant from a healthy donor. Data was collected from 2 independent experiment and is shown as median with interquartile range. B) Flow cytometry analysis of CD69 and eotaxin-1 expression in MAIT cells. C) Flow cytometry analysis of CD69 and granzyme A expression in eosinophils. (B and C) Expression of each marker is shown as MFI (median fluorescence intensity). Each dot represents a sample from an individual healthy donor. Data was collected from 4 independent experiments. **p<0.01 *** p<0.001 **** p<0.0001 by Wilcoxon ranked test.

Previous studies have demonstrated the ability of eosinophils to kill COLO205 cells in a granzyme A-dependent manner (43). To determine if MAIT cell activation can induce eosinophil activation and cytotoxic function in a setting with mixed cell population of leukocytes, we cultured human peripheral leukocytes with or without the MAIT ligand 5-A-RU/MGO overnight and analyzed cytokine expression and levels of activation of MAIT cell and eosinophil using flow cytometry (Supplemental Fig. 1). Ligand stimulated MAIT cells had significantly higher expression of the activation marker CD69 (p<0.01) and of eotaxin-1 (p<0.001) (Fig. 4B). Compared to unstimulated cultures, whole leukocyte cultures stimulated with the MAIT ligand also shows a significant increase in eosinophil activation, marked by CD69 and granzyme A expression (p<0.0001; Figure 4C). These data show that in peripheral leukocyte populations, the addition of the MAIT ligand induces the expression of eosinophil-activating cytokines in MAIT cells and correspondingly, eosinophil activation.

To confirm if eosinophil activation upon 5-A-RU/MGO stimulation is mediated via MR1 or a broad consequence of antigen processing, anti-MR1 neutralizing antibody was added to the human leukocyte cultures 1 hour before stimulation with 5-A-RU/MGO. In the presence of anti-MR1, MAIT cell activation and the expression of eotaxin-1 are significant reduced (p < 0.001 and p < 0.0001; Fig. 4B). Similarly, the activation and the induction of effector molecule granzyme A in eosinophils induced by MAIT cell stimulation are significantly reduced in the presence of anti-MR1 antibody (p < 0.001; Fig. 4C), suggesting that the effect of 5-A-RU/MGO on eosinophils is dependent on MR1.

Together, these data show that the addition of 5-A-RU/MGO, a MAIT-specific ligand, leads to an increased production of eosinophil-recruiting chemokines and cytokines by MAIT cells and eosinophil activation, highlighting the ability of MAIT cells to modulate eosinophils in a MR1-dependent manner.

## Discussion

While MAIT cells are primarily known for their ability to kill microbially infected cells, other potential roles have emerged. Specifically, due to their abundance at mucosal tissues, there has been an increasing interest in the role of MAIT cells in mucosal tissue-associated cancers. MAIT cells are a promising potential candidate for universal cancer immunotherapies: MAIT cells lack alloreactive potential, thus having an advantage over the use of conventional T cells with HLA-restrictions for the application of non-donor-restrictive adoptive cell therapy (24). MAIT cells also express higher levels of the multi-drug efflux protein MDR1, conferring higher persistence against certain chemotherapy drugs compared to conventional CD8^+^ T cells and thus an ideal target for use in immunotherapy coupled with chemotherapy (25, 26, 44). Additionally, the checkpoint inhibitor, anti-PD1, has been shown to enhance the cytotoxicity of MAIT cells in patients with prostate cancer (45), basal and squamous cell carcinoma, and hepatocellular carcinoma (46–48). Recent studies have demonstrated the ability to generate chimeric antigen receptor (CAR)-MAIT cells, which retain their anti-tumor capacity even in the presence of immunosuppressive tumor-associated macrophages (49, 50). Furthermore, protocols for MAIT cell expansion have been established, including demonstration that, when expanded with a human serum replacement, MAIT cells display enhanced functionality and low levels of exhaustion markers, making them optimized for adoptive therapy (51, 52).

In this study, we provided evidence that MAIT cells have an anti-tumoral role both in an MC38 cell-derived allograft tumor model in RAG1^-/-^ mice and against human colon cancer cells. We used RAG deficient mice which are characterized by the lack of mature B and T cells to exclude the potential impact of the presence of endogenous T cells. In the murine model of colon cancer, we observed tumor inhibition, associated with a significant increase in the production of anti-tumor cytokines, and evidence suggesting the recruitment of eosinophils with increases in both gene and protein expression of eotaxin, IL-5, and SiglecF, among others. In human leukocyte and human colon cancer cell co-culture, we observed enhanced levels of COLO 205 lysis after stimulation with MAIT ligand 5-A-RU/MGO, demonstrating their cytotoxic capacity against colon cancer cells. In addition, MAIT cell activation in human whole leukocytes leads to the production of a wide range of cytokines and chemokines, including eotaxin-1, GM-CSF, and Mip-1, highlighting the potential for anti-tumor effects through their immune-modulating properties (10). Particularly, we observed that MAIT cell activation resulted in an increase in the production of IL-5 and eotaxin-1, and GM-CSF, and an increase in activation and effector function of eosinophils, the effect of which was MR1-dependent.

We observed the capacity of MAIT cell to modulate innate cell populations in our models of colon cancer. We found significantly higher levels of eotaxin-1, known to recruit and activate eosinophils, in both mouse *in vivo* and human *in vitro* models. *In vitro* studies have demonstrated that eosinophils have the ability to induce direct killing on colon cancer cells (43, 53), supported by an *in vivo* study of two colon cancer models showing that eosinophil contributes to tumor suppression (53). Another study using a subcutaneous MC38 cells-derived tumor model in WT mice has shown that eosinophil inversely correlates with tumor growth and the observed tumor suppression was dependent on GM-CSF and eosinophil recruitment (54). This is consistent with our experiment where we observed an association between MAIT cell-mediated inhibition of MC38-derived tumors and an increase in tumor production of GM-CSF and eotaxin-1. With eosinophils’ known ability to release effector molecules like major basic protein (MBP) into the tumor microenvironment to induce cancer cell lysis (55), these data support the involvement of eosinophils in MAIT-mediated tumor suppression.

In addition to the production of eosinophil modulating effectors, we report MAIT cell production of other pro-inflammatory and immune-modulating cytokines, GM-CSF, IFN-γ, and IL-17. GM-CSF regulates the proliferation, differentiation, recruitment, and effector functions of myeloid cell populations (11, 12, 56), and its addition of GM-CSF to human primary monocytes inhibits COLO 205 proliferation (57). In murine colon cancer, neutrophils play a role in limiting tumor progression, invasion and proliferation (58). We found higher GM-CSF in tumors from the MAIT cell-injection group, suggesting a potential role for myeloid cell populations in mediating the observed anti-tumor effect. We also found increased expression of IFN-γ in MAIT-treated tumors. IFN-γ is a cytotoxic cytokine that can act directly on tumor cells to induce tumor elimination (59–62). Specifically, IFN-γ inhibits MC38-derived tumor growth in mice (63) and COLO 205 in vitro (64). In addition to the direct effects on tumor cells, IFN-γ also has immune modulating effects on innate immune cells, including the enhancement of the anti-tumor activity of eosinophils against colon cancer cell lines, including MC38 (53). Finally, our demonstration of increased IL-13 and IL-17 in MAIT-treated tumors is consistent with studies showing their production by MAIT cells (26, 65, 66). In line with our murine model of colon cancer, a prior study showed that treatment with recombinant IL-17 in MC38-bearing mice results in tumor inhibition (63).

Taken together, our study employing both murine model and human cell cultures demonstrates a protective role of MAIT cells in colon cancer. While further analysis of immune cell activity in the tumor microenvironment is needed to fully understand the interaction between MAIT cells and other immune cells, our findings suggest the potential for MAIT cells in recruitment and activation of eosinophils. The mechanism by which MAIT cell mediates MC38-derived tumor inhibition, particularly the role of eosinophils, needs further study, including how MAIT cells could be harnessed in future therapeutic applications against colon cancer.

## Methods and Materials

### Mouse and cell lines

CL57BL/6 wild type and B6.129S7-*Rag1^tm1Mom^*/*J* (*Rag1*^−/−^) mice were purchased from the Jackson Laboratory (Bar Harbor, MA, USA). Mice were housed at the University of Utah CMC Animal Facility on a 12h – 12h light-dark cycle and were provided with water and normal rodent chow ad libitum. Mouse and human cell lines, MC38 and Colo205, were purchased from the American Type Culture Collection (ATCC, Manassas, Virginia, United States) and maintained in RPMI complete media supplemented with 10% fetal bovine serum (FBS, Thermo Fisher Scientific, Waltham, MA, USA), 1% penicillin/streptomycin cocktail (Corning, Tewksbury, MA, USA) and 25mM HEPES (Thermo Fisher Scientific, Waltham, MA, USA).

### *In vivo* MAIT cell expansion and isolation

On day 0, C57BL/6J mice were anesthetized with isoflurane and intranasally inoculated with a 50 uL solution (25 uL per nare) of PBS containing 200 nmol 5-A-RU (67), 50 mM methylglyoxal (Sigma-Aldrich, St. Louis, MO, USA), and 16.67 mg Pam2CSK4 (InvivoGen, San Diego, CA, USA). The reaction of methylglyoxal with 5-A-RU results in the formation of the MAIT-activating ligand 5-(2-oxopropylideneamino)-6-D-ribitylaminouracil (5-OP-RU) (68). On days 1, 2 and 4, mice were again inoculated with 200 nmol 5-A-RU and 50 mM methylglyoxal. On day 7, mice were euthanized, lungs were perfused with 5 mL of PBS, and then single-cell lung suspensions were prepared using the gentleMACS Mouse Lung Dissociation Kit (Miltenyi Biotech, Bergisch Gladbach, North Rhine-Westphalia, Germany) according to the manufacturer’s protocol. Lung suspensions were then pooled into male and female groups and stained with fluorochrome-conjugated antibodies for MAIT isolation using fluorescent-associated cell sorting (FACS). Lung suspensions were first incubated with Fixable Viability Dye eFluor 780 (BD Biosciences, Franklin Lakes, NJ, USA) and then blocked with anti-mouse CD16/CD32 Fc block (Biolegend, San Diego, CA, USA) and unlabeled MR1-6-formylpterin-tetramer (6-FP) (NIH Tetramer Core Facility at Emory University, Atlanta, GA, USA) to minimize non-specific binding. Suspensions were then stained for 30 minutes at room temperature with the MAIT cell TCR binding MR1-5-OP-RU-tetramer (NIH Tetramer Core Facility at Emory University, Atlanta, GA, USA) and the following antibodies: anti-CD3-PE-Dazzle594, anti-TCRb-BV421, anti-CD4-FITC, anti-CD8-APC, anti-TCRgd-PE-Cy7, anti-CD44-BV650, and anti-CD45R-PE-Cy5 (Supp. Table 1). MAIT cells were gated as single cell lymphocyte, live, B220-, TCR γδ-, CD44^hi^, CD3+, TCRβ+ and MR1-5OPRU+. Following FACS, sorted MAIT cells were washed in sterile PBS and resuspended at 1x10^6^ cells per ml for use for peritumoral injection in the MC38 cancer model.

### MAIT cell peritumoral injection in MC38 mouse model

MC38 cell inoculation and MAIT cell peritumoral injection were performed as previously described (69). Briefly, in Rag1-/- mice, 1x10^6^ MC38 cells were injected subcutaneously on Day 0. 1x10^5^ FACS-sorted, sex-matched MAIT cells were then injected peritumorally (same site as MC38 injection) on Day 1. For control group, mice were injected with an equivalent volume of PBS. Tumor growth was monitored by measuring tumor size with caliper until Day 15-21 and the mice were sacrificed to obtain the tumors. Extracted tumors were weighed at the end of the experiment.

### Mouse MAIT cell in vitro stimulation

Expanded lung MAIT cells from C57BL/6J donor mice were cultured in vitro at 50,000 cells/well. Cells were either left unstimulated or stimulated with 1X eBioscience Cell Stimulation Cocktail (500X, Invitrogen, Thermo Fisher Scientific, Waltham, MA, USA) in 100uL complete RPMI (Gibco, Thermo Fisher Scientific, Waltham, MA, USA, supplemented with 10% FBS, 1% penicillin/streptomycin, 1mM HEPES). After overnight culture, supernatants were analyzed using MILLIPLEX MAP Mouse Cytokine/Chemokine Magnetic Bead Panel - Premixed 25 Plex - Immunology Multiplex Assay (EMD Millipore Corporation, Billerica, MA, USA). and custom FAS panel on the Luminex MAGPIX. A complete list of cytokines and chemokines analyzed is listed in Supplemental Table 1.

### Detection of tumor-released cytokines

8 mg (± 0.5mg) of each mouse tumor was incubated in 500 µL of complete RMPI medium for 18 hours and 25 µL of the resulting supernatant was analyzed using Luminex bead assay for various soluble markers using MILLIPLEX MAP Mouse Cytokine/Chemokine Magnetic Bead Panel - Premixed 25 Plex - Immunology Multiplex Assay (EMD Millipore Corporation, Billerica, MA, USA). and custom FAS panel on the Luminex MAGPIX. A complete list of cytokines and chemokines analyzed is listed in Supplemental Table 1. Measurements below detectable limits were considered as 0.

### Caspase activity measurement

Apo-ONE® Homogeneous Caspase-3/7 Assay (Promega Corporation, Madison, WI, USA) was used to determine the caspase activity in the extracted tumors. 8 mg tumor pieces were homogenized in Apo-ONE® Caspase-3/7 Reagent using a bead beater. After 1 freeze-thaw to ensure lysis, samples were incubated for 30 minutes on a shaker and analyzed according to manufacturer’s protocol. Caspase 3/7 activity is measured as arbitrary fluorescent unit (AFU) using a SpectraMax Mini plate reader (Molecular Devices, San Jose, CA, USA).

### Human whole leukocyte preparation and culture

Human whole leukocytes from anonymous healthy donors were isolated from apheresis cones from ARUP using RBC sedimentation with HetaSep (STEMCELL Technologies, Vancouver, British Columbia). Freshly isolated whole leukocytes were used fresh for *in vitro* culture.

Whole leukocytes were cultured with or without 5-A-RU (50 nM) and MGO (50 µM) in 200 µL RPMI complete media and incubated at 37 °C overnight. CD69, and eotaxin, granzyme A, and in MAIT cells and Eosinophils were analyzed using flow cytometry. To detect released cytokines in the medium, 25 µL of supernatant was analyzed using Cytokine 35-Plex Human Panel for the Luminex™ platform (Invitrogen, Thermo Fisher Scientific, Waltham, MA, USA) on the Luminex MAGPIX. A complete list of cytokines and chemokines analyzed is listed in Supplemental Table 2. To block interactions activation of MAIT cell via MR1, anti-MR1 (Clone 26.5, Biolegend, San Diego, CA, USA) or an isotype control was used at 10 µg/ml.

### COLO 205 Killing Assays

For Calcein Release Assay, 50,000 Calcein-labeled (2 µM, Biolegend, San Diego, CA, USA) COLO 205 cells were co-cultured with 500,000 human whole leukocytes for 4 hours with or without 5-A-RU (50 nM) and MGO (50 µM) in 200 µL RPMI complete media at 37 °C. Maximum release was set up with 2% Triton X-100 (Acros Organics, Thermo Fisher Scientific, Waltham, MA, USA). All samples and controls were set up in triplicates. Cultures were gently mixed and centrifuged, and 100 µL of culture supernatant was transferred to a new plate and analyzed for fluorescence intensity using BioTek Synergy H1 Multimode Reader (BioTek Instruments, Inc, Winooski, VT, USA). Percent Specific Lysis was expressed as a percent relative to maximum release control, calculated as [(Fluorescence _Sample_ – Fluorescence _Untreated_)/ (Fluorescence _max_ – Fluorescence _Untreated_)] x 100. For flow cytometry-based killing assay, 100,000 CFSE-stained (Immunochemistry Technologies, Davis, California, CA, USA) COLO 205 was co-cultured with 10^6^ human whole leukocytes for 16 hours and the frequency of CFSE+ cells was analyzed.

### Flow Cytometry and FACS

Isolated mouse lung cells were stained and MAIT cells were sorted using FACS Aria III Cell Sorter (BD Bioscience). Stained human *in vitro* samples were analyzed using Cytek Aurora (Cytek Biosciences, Fremont, CA, USA). FCS files were analyzed using FlowJo^TM^ v10.8 Software (BD Life Sciences, Franklin Lakes, NJ, USA). MAIT cells were defined from the lymphocyte as live singlet, CD45^+^, CD3^+^, MR1-5-OP-RU-Tetramer^+^, Vα7.2^+^ cells. Eosinophils were gated from the granulocyte population as live singlet, CD45^+^, CD3^-^, CD44^+^, Siglec8^+^ cells. Gating strategies are included in Supplemental Figure 1.

### Antibodies used for FACS and Flow Cytometry

List of antibodies used in FACS of mouse MAIT cells.

| Antibody/Viability dye | Clone | Vendor |
| --- | --- | --- |
| eBioscience Fixable<br>Viability Dye eFluor™<br>780 | NA | Invitrogen, Thermo Fisher<br>Scientific |
| Anti-CD45R/B220 | RA3-6B2 | Biolegend |
| Anti-gd | GL3 | Biolegend |
| Anti-CD3 | 17A2 | Biolegend |
| Anti-CD44 | IM7 | Biolegend |
| MR1-5-OP-RU | NA | NIH Tetramer Core |
| Anti-TCRb | H57-597 | Biolegend |

List of antibodies used in flow cytometry analysis of human leukocyte cultures.

| Antibody/Tetramer | Clone | Vendor |
| --- | --- | --- |
| Zombie UV | NA | Biolegend |
| Anti-CD45 | HI30 | Biolegend |
| Anti-CD45RO | UCHL1 | BD Biosciences |
| Anti-CD3 | SK7 | BD Biosciences |
| Anti-CD4 | SK3 | BD Biosciences |
| Anti-CD8 | RPA-T8 | Biolegend |
| MR1-5OPRU | NA | NIH Tetramer Core |
| Anti-Vα7.2 | 3C10 | Biolegend |
| Anti-CD161 | HP-3G10 | Biolegend |
| Anti-CD25 | BC96 | Biolegend |
| Anti-CD69 | FN50 | BD Biosciences |
| Anti-CD44 | IM7 | Biolegend |
| Anti-Siglec8 | 7C9 | Biolegend |
| Anti-Granzyme A | CB9 | Biolegend |
| Anti-Eotaxin-1 | IC3201U | R&D Systems |

### Statistical Analysis

Statistical analysis was performed using GraphPad Prism version 8 for Mac (GraphPad Software, Boston, Massachusetts USA). Two-tailed Mann-Whitney U test was used to compare tumor weights between groups, and Wilcoxon ranked t test was used to compare variables between treatment groups in experiment involving paired human samples.

## Author Contributions

OJC, EJB, and DTL designed the experiments. OJC, EJL, OJ, DJ, and ST performed the experiments. OJC analyzed the data. EJB and DTL supervised the work. OJC, OJ, EJB and DTL wrote the paper. EJB and DTL obtained funding. All authors reviewed and approved the final manuscript.

## Acknowledgement

The research was supported by the National Institutes of Health (R01CA207051 to EJB, R01AI130378 to DTL, and K24AI166087 to DTL). We would like to thank the University of Utah Flow Cytometry Core.

