## Supplemental Figures 1-3; Supplemental Tables 1-2 for "MAIT Cells Modulate Innate Immune Cells and Inhibit Colon Cancer Growth"

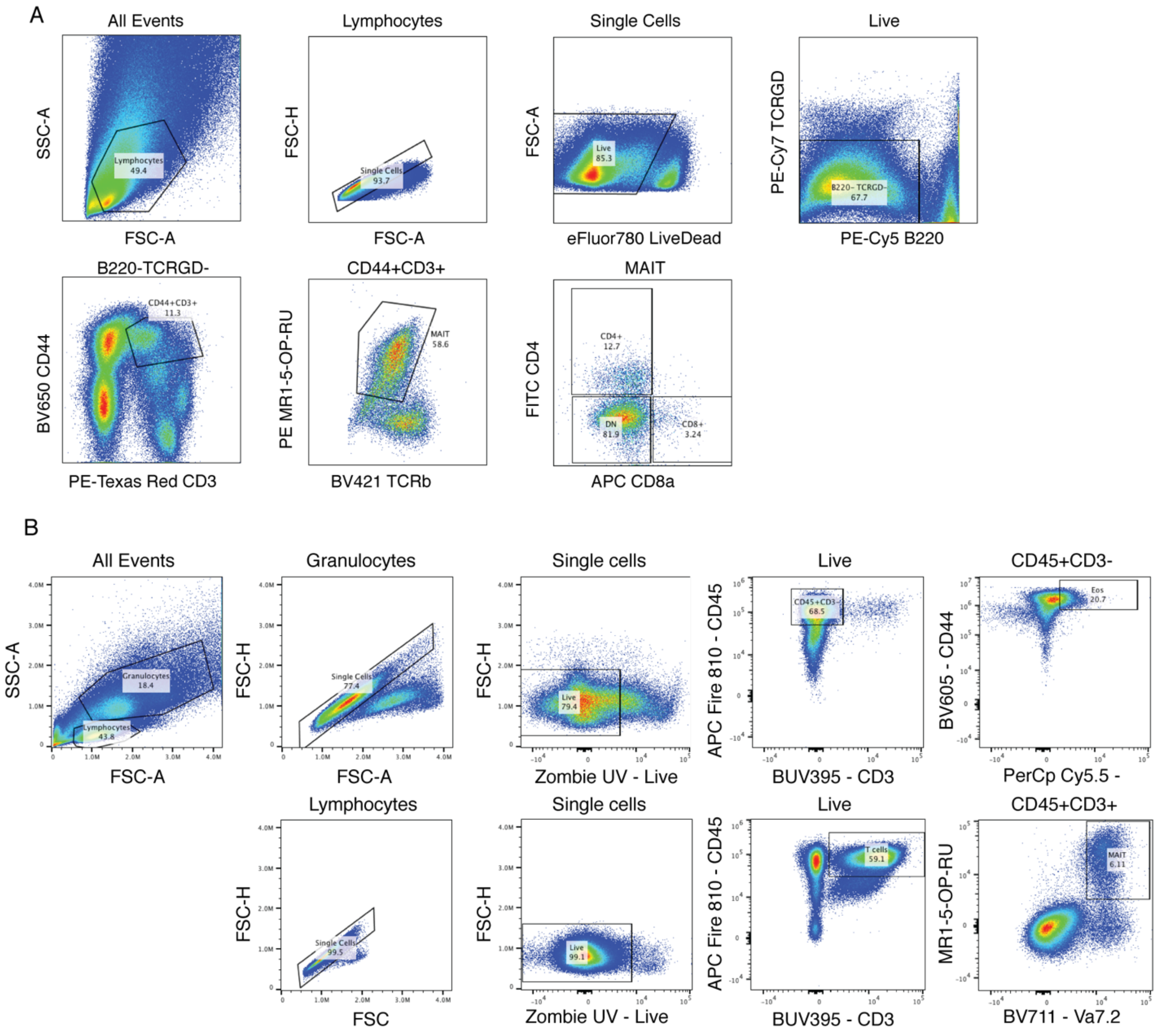


**Supplemental Figure 1. FACS sorting strategy for the purification of MAIT cells expanded in C57BL/6J mice and flow cytometry gating strategy of human MAIT cells and eosinophils within whole leukocyte cultures.**

A) MAIT cells expanded *in vivo* were sorted from dissociated lung suspension using FACS Aria III Cell Sorter. MAIT cells were gated as single cell lymphocyte, live, B220$^-$, TCR$\gamma\delta^-$, CD44$^{hi}$, CD3$^+$, TCR$\beta^+$ and MR1-5OPRU$^+$. B) Representative flow plot showing the gating strategy to identify eosinophils (Granulocyte, single cell, live, CD45$^+$, CD3$^-$, CD44$^+$, Siglec8$^+$) and MAIT cell (Lymphocyte, single cell, live, CD45$^+$, CD3$^+$, MR1-5-OP-RU$^+$, V$\alpha$7.2$^+$).

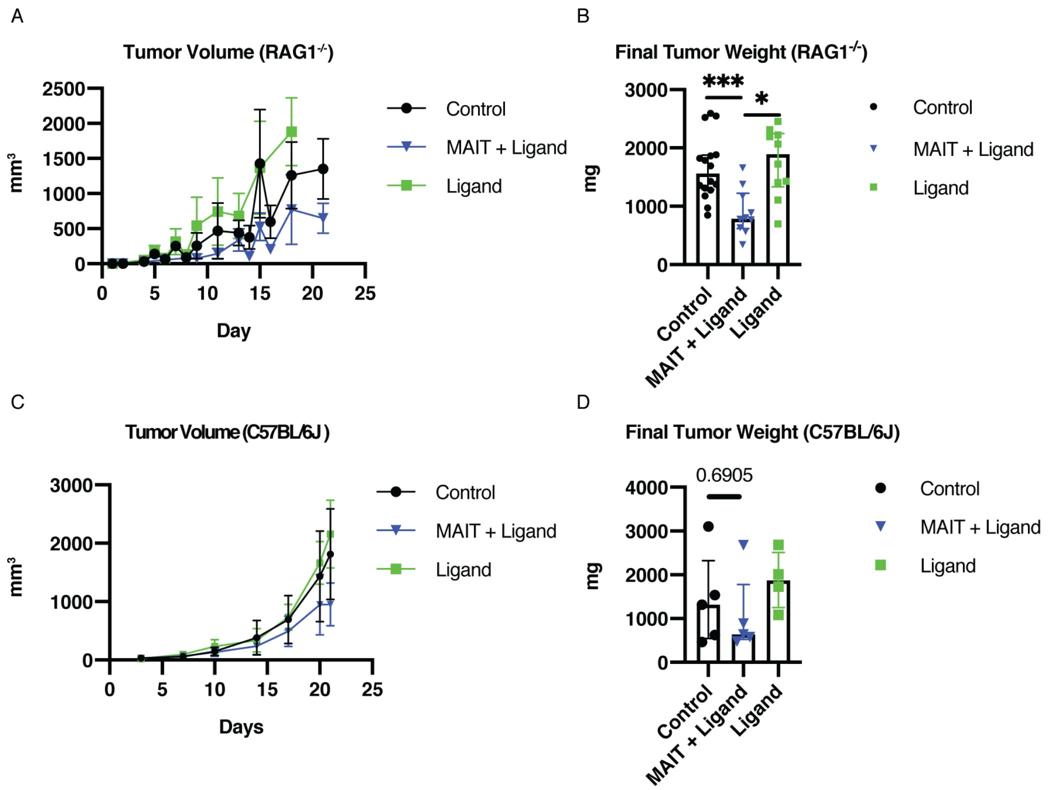


**Supplemental Figure 2. Peritumoral injection of MAIT cell and ligand inhibits tumor growth in RAG1$^{-/-}$ and C57BL/6J mice.**

A) MC38 cells-derived tumor volume in mm$^3$ over time in RAG1$^{-/-}$ mice with MAIT cell and ligand injection compared to PBS control. Control: n = 16 (11 females and 5 males). MAIT + Ligand: n = 10 (5 females and 5 males). Ligand: n = 10 (5 females and 5 males). B) Weight of extracted MC38 cells-derived tumors from RAG1$^{-/-}$ mice with or without MAIT cell and ligand injection in mg. C) MC38 cells-derived tumor volume in mm$^3$ over time in C57BL/6J mice with MAIT cell and ligand injection compared to PBS control. N = 5 female mice for all three groups. D) Weight of extracted MC38 cells-derived tumors from C57BL/6J mice with or without MAIT cell and ligand injection in mg. Each dot represents a tumor sample and data is shown as median with interquartile range. *p<0.05 *** p<0.001 by two-tailed Mann-Whitney U test.

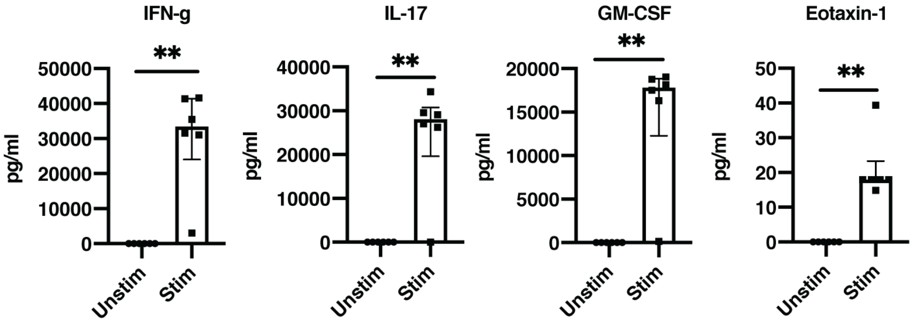


**Supplemental Figure 3. Sorted mouse MAIT cells produce a wide variety of cytokines and chemokines *in vitro*.**

Concentration of various cytokines and chemokines from culture of sorted MAIT cells. 100,000 MAIT cells were cultured overnight with or without 1X T cell activation cocktail. Supernatant was analyzed using Luminex multiplex assay. Each dot represents a replicate and data is shown as median with interquartile range. **p<0.01 by two-tailed Mann-Whitney U test.

| | | | | |
|---|---|---|---|---|
| g-csf | eotaxin | gm-csf | ifng | il-1a |
| il-1b | il-2 | il-4 | il-3 | il-5 |
| il-6 | il-7 | il-9 | il-10 | il-12p40 |
| il-12p70 | lif | il-13 | lix | il-15 |
| il-17 | ip-10 | kc | mcp-1 | mip-1a |
| m-csf | mip-2 | mig | rantes | vegf |
| tnfa | | | | |

**Supplemental Table 1. Complete list of mouse cytokines and chemokines analyzed with the Mouse Luminex Cytokine Panel.**

| | | | | |
|---|---|---|---|---|
| EGF | Eotaxin | TGF-a | G-CSF | FLT-3L |
| GM-CSF | Fractalkine | IFNa2 | IFNy | GRO |
| L-10 | MCP-3 | IL-12p40 | MDC | IL-12p70 |
| PDGF-AA | IL-13 | PDGF-AB/BB | IL-15 | sCD40L |
| IL-17A | IL-1RA | IL-1a | IL-9 | IL-1B |
| IL-2 | IL-3 | IL-4 | IL-5 | IL-6 |
| IL-7 | IL-8 | IP-10 | MCP-1 | Mip-1a |
| Mip-1B | RANTES | TNFa | TNFb | VEGF |

**Supplemental Table 2. Complete list of human cytokines and chemokines analyzed with the Human Luminex Cytokine Panel**
